# Characterization of hormone-producing cell types in the teleost pituitary gland using single-cell RNA-seq

**DOI:** 10.1101/2020.12.14.422690

**Authors:** Khadeeja Siddique, Eirill Ager-Wick, Romain Fontaine, Finn-Arne Weltzien, Christiaan V. Henkel

**Affiliations:** Physiology Unit, Faculty of Veterinary Medicine, Norwegian University of Life Sciences, Ås, Norway

## Abstract

The pituitary is the vertebrate endocrine gland responsible for the production and secretion of several essential peptide hormones. These, in turn, control many aspects of an animal’s physiology and development, including growth, reproduction, homeostasis, metabolism and stress responses. In teleost fish, each hormone is presumably produced by a specific cell type. However, key details on the regulation of, and communication between these cell types remain to be resolved. We have therefore used single-cell sequencing to generate gene expression profiles for 2592 and 3804 individual cells from the pituitaries of female and male adult medaka (*Oryzias latipes*), respectively. Based on expression profile clustering, we define 15 and 16 distinct cell types in the female and male pituitary, respectively, of which ten are involved in the production of a single peptide hormone. Collectively, our data provide a high-quality reference for studies on pituitary biology and the regulation of hormone production, both in fish and in vertebrates in general.

## Background & Summary

The pituitary is a master endocrine gland in vertebrates, which is involved in the control of a variety of essential physiological functions including growth, metabolism, homeostasis, reproduction, and response to stress^1–3^. These functions are modulated by the secretion of several peptide hormones, produced by different endocrine cell types of the adenohypophysis^2^.

While in mammals the endocrine cells are distributed throughout the adenohypophysis^2,4^, the pituitary of teleost fish is highly compartmentalized, with specialized hormone-producing cell types located in specific regions^2^. As the teleost pituitary produces at least eight peptide hormones in distinct cell types^2^, the physiology of the pituitary gland is relatively complex^5^. The *rostral pars distalis* contains lactotropes and corticotropes, which produce prolactin (Prl) and adrenocorticotropic hormone (Acth), respectively. Somatotropes (producing growth hormone, Gh), gonadotropes (luteinizing hormone, Lh and follicle-stimulating hormone, Fsh) and thyrotropes (thyroid-stimulating hormone, Tsh) are located in the *proximal pars distalis*, and somatolactotropes (somatolactin, Sl) and melanotropes (α-melanocyte stimulating hormone, α-Msh) in the *pars intermedia*^4^. In addition, in tetrapods, the gonadotropins Lh and Fsh can be produced in the same cell, while fish gonadotropes are generally thought to secrete either Fsh or Lh, but not both^2,6–8^.

In recent years, transcriptome sequencing (RNA-seq) has emerged as a powerful technology to study the expression and regulation of pituitary genes. For example, RNA-seq has been used to study gene expression in the zebrafish pituitary during maturation^9^, in prepubertal female silver European eel pituitary glands^10^, and in FACS-selected Lh cells from the Japanese medaka pituitary^11^. Although these studies provide a valuable perspective on the regulation of hormone production, RNA-seq is a bulk technology, which averages gene expression over the entire tissue sample. As a result, it does not provide information on which hormones are produced in which cells. Knowledge on the physiology and development of the individual pituitary cell types, as well as on the regulatory mechanisms involved in hormone production, therefore remains limited.

In this study, we employed single-cell RNA-seq (scRNA-seq) technology, which allows the study of biological questions in which cell-specific changes in the transcriptome are important, e.g. cell type identification, variability in gene expression and heterogeneity within populations of genetically identical cells^12,13^. scRNA-seq has recently been used to describe the heterogeneity within the pituitary cell populations in mammals^14,15^ and zebrafish^16^. Unlike bulk RNA-seq, scRNA-seq promises to disentangle the processes at work in pituitary hormone production.

We used medaka (*Oryzias latipes*)^17,18^ to construct a transcriptomic snapshot of the cell types in the adult pituitary gland. Medaka is an emerging model species, which has a small genome (800 Mbp) and is a representative of the largest radiation within the teleosts (euteleosts, Cohort^19^ Euteleostomorpha). As such, it is a complementary model to zebrafish^16^, which belongs to another major teleost clade (Cohort Otomorpha)^19^.

We used the 10x Genomics scRNA-seq platform to analyze female and male pituitaries separately and obtained 2592 and 3804 high-quality cellular transcriptome profiles, respectively. This dataset provides a comprehensive transcriptomic reference on the teleost pituitary, as well as on its constituent cell types. After bioinformatics analysis we found these profiles belonging to 15 and 16 distinct cell populations in the female and male pituitary, respectively, of which ten are involved in the production of a single peptide hormone. This suggests the teleost pituitary is subject to a higher degree of division of labour than its mammalian counterpart, in which many cells express multiple hormone-encoding genes^14^.

## Methods

### Fish strain and animal husbandry

For this study, we used a transgenic line of Japanese medaka derived from the d-rR genetic background, in which the Lhβ gene (*lhb*) promoter drives the expression of the green fluorescent protein (*hr-gfpII*) gene^20,21^. Fish were kept in a re-circulating system (up to 10–12 fish per 3-liter tank) at 28 °C on a 14/10h light/darkness cycle and were fed three times a day with a mix of dry food and Artemia. The animal experiments performed in this study were approved by the Norwegian University of Life Sciences, following guidelines for the care and welfare of research animals.

### Pituitary sampling and cell dissociation

Pituitaries were collected from 24 female and 23 male adult medaka of approximately 8 months old (eggs fertilized 20 October 2017, sampling 6 June 2018). At this age, medaka are fully mature and sexually dimorphic. Fish were sacrificed between 07.30–09.00h in the morning (around the time of spawning induced by the start of the artificial light period) by immersion in ice water followed by severing of the spinal cord, and immediate dissection of the pituitary. In order to reduce the influence of sampling time on the results, fish were processed in batches of 10 individuals of each sex at a time.

After sampling, pituitaries cells were dissociated as described^22^. Briefly, fresh pituitaries were put in an Eppendorf tube with modified PBS (phosphate buffered saline, pH 7.75, 290 mOsm/kg, 0.05% bovine serum albumin) and kept on ice until processing. Pituitaries were treated with 0.1% w/v trypsin type II S for 30 minutes at 26 °C, then with 0.1% w/v trypsin inhibitor type I S for 20 minutes at 26 °C, and subsequently mechanically dissociated using a glass pipette. Clumps and non-cellular debris were removed by filtration. Dissociated pituitary cells were resuspended in 90 μl modified PBS and kept on ice until scRNA-seq library preparation, which commenced at 12.00h on the same day.

### 10x Genomics library preparation

We prepared scRNA-seq libraries on the 10x Genomics Chromium Controller at the Genomics Core Facility of the Radium Hospital (part of Oslo University Hospital). For the initial step of the library preparation (GEM generation and barcoding) 35 μl of the cell suspension prepared in the previous step was used, containing approximately 10,000 (female) and 11,000 (male) cells. The 10x Genomics Single Cell 3’ Reagents Kits v2 were used according to the manufacturer’s guidelines^23^ to prepare cDNA libraries for a target of 4,000 cells.

### Sequencing

Per sample, four redundant sequencing libraries were analyzed on an Illumina NextSeq 500 at the Genomics Core Facility. Read 1 was sequenced for 28 cycles (covering the 26 nt cellular barcode and UMI), read 2 for 96 cycles (covering the cDNA insert). For downstream analyses, the resulting FASTQ files were collated per sample.

### Reference genome and annotation

We used the chromosome-scale Hd-rR medaka reference genome (Ensembl release 94) for alignments. In preliminary analyses, we noticed that many reads for expected pituitary-specific transcripts were mapped near, but outside the 3’ boundary of gene annotations. Therefore, in order to improve transcript quantification accuracy, we manually updated the annotation of these and other highly expressed genes (Table 1). The resulting custom gene transfer file (GTF) was then supplied using the 10x Genomics Cell Ranger^24^ (*v 3.0.2) mkref* command.

**Table 1.**
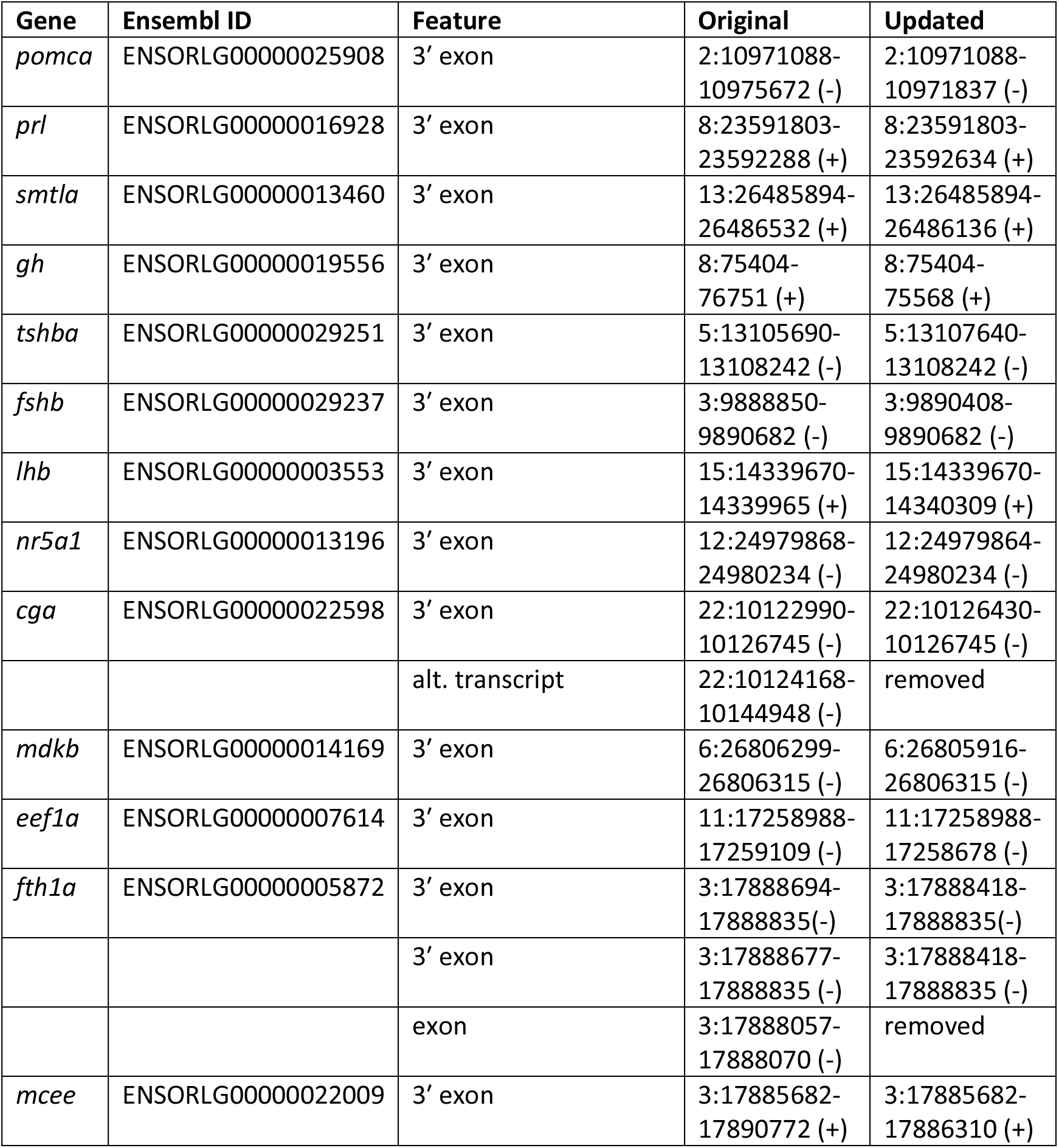
Genome annotation adjustments.

### Initial data processing

We used the standard 10x Genomics Cell Ranger^24^ (*v 3.0.2*) pipeline with default parameters for preliminary analyses. It includes quality control of the FASTQ files, alignment of FASTQ files to the customized medaka reference using STAR^25^ (*v 2.5.1b*), demultiplexing of cellular barcodes, and quantification of gene expression. Table 2 summarizes the Cell Ranger quality control and output.

**Table 1.** Sequencing and Cell Ranger statistics.

|  | Female | Male |
| --- | --- | --- |
| Number of reads | 165818560 | 142738097 |
| Q30 bases in barcode | 97.5% | 97.4% |
| Q30 bases in RNA read | 79.7% | 80.3% |
| Q30 bases in UMI reads | 83.4% | 81.5% |
| Reads mapped to genome | 84.1% | 85.4% |
| Reads mapped confidently to exonic regions | 59.9% | 62.5% |
| Reads mapped uniquely to genome | 82.4% | 83.8% |
| Estimated number of cells | 2890 | 4321 |
| Fraction of reads in cells | 83.4% | 81.5% |
| Mean reads per cell | 57376 | 33033 |
| Median genes detected per cell | 1182 | 1063 |
| Total genes detected | 17321 | 17775 |

Cell Ranger also performs automated cell clustering based on the similarity of gene expression profiles. We used the Loupe Cell Browser^26^ (*v 3.1.1*) to visualize these results. Unsupervised graph-based clustering suggested nine and eleven cell populations in the female and male medaka pituitary, respectively (see Code Availability).

For a more detailed analysis, we used the Seurat^27^ R toolkit (*v 3.1.5*) and dropEst ^28^ (*v 0.8.5*) for accurate estimation of molecular counts per cell (Table 3). We used the .bam files produced by Cell Ranger as the input for dropEst.

**Table 3.** Final cell quantification statistics.

|  | Female | Male |
| --- | --- | --- |
| Unfiltered cells | 74338 | 77520 |
| Filtered cells | 4945 | 7334 |
| Filtered genes detected | 18786 | 19043 |
| Genes with exons | 17284 | 17882 |
| Genes with introns | 15036 | 15598 |
| Cells pre-doublet detection | 2644 | 3921 |
| Cells post-doublet detection | 2592 | 3804 |

### Quality control (QC)

dropEst by default reports any barcode with more than 100 UMIs as cellular (filtered cells in Table 3). Presumably, this relaxed threshold still includes many non-cellular and debris barcodes. Therefore, we explored these data using Seurat, and derived additional selection criteria for cellular barcodes based on the UMIs counts, mitochondrial fraction and globin expression. After imposing these thresholds, 2644 and 3921 cells remained for the female and male pituitary, respectively.

Since single-cell capture is sensitive to the formation of doublets, we used the R package DoubletFinder^29^ (*v 2.0.3*) to detect hybrid expression profiles in our data. The number of expected real doublets depends on the number of cells captured. We selected a 2% doublet rate for the female sample and a 3% doublet rate for the male sample, based on the 10x Genomics specifications^30^. We removed 52 and 117 putative doublets from the female and male pituitary data, respectively.

### Clustering

After QC, we normalized the data with the *LogNormalize* method in Seurat and identified highly variable genes by using the *vst* method. Subsequently, we used these variable genes (*n* = 2000) to identify significant principal components (PCs) based on the *jackStraw* function. We used ten informative PC dimensions in both samples as the input for uniform manifold approximation and projection (UMAP). After the dimensionality reduction, we clustered the cells using the *FindClusters* function with a resolution of 0.5 and 0.9 for the female and male data sets, respectively, and initially characterized 11 different clusters in both the female and male pituitary.

### Cell type assignment

We used differentially expressed genes per cluster (*FindAllMarkers* with *min.pct = 0.25*) to help assign tentative cell type identities to these clusters. Marker genes (Table 4) reported in previous studies^2,14–16,21,31,32^ were used for cell type assignment.

**Table 4.** Cells per cluster in scRNA-seq data.

| Cell types | Hormone | Female cells | % | Male cells | % | Marker gene | References |
| --- | --- | --- | --- | --- | --- | --- | --- |
| 1. Melanotropes | α-Msh | 54 | 2.1 | 53 | 1.4 | *Pomca, oacyl* | <sup>2,15</sup> |
| 2. Corticotropes | Acth | 24 | 0.9 | 55 | 1.5 | *pomca* | <sup>15</sup> |
| 3. Lactotropes | Prl | 148 | 5.7 | 175 | 4.6 | *prl* | <sup>14,15</sup> |
| 4. Lactotropes | Prl | 66 | 2.6 | 222 | 5.8 | *prl* | <sup>14,15</sup> |
| 5. Somatolactotropes | Sl | 15 | 0.6 | 38 | 0.9 | *smtla* | <sup>16,31</sup> |
| 6. Somatotropes | Gh | 53 | 2.0 | 127 | 3.3 | *gh* | <sup>14,15</sup> |
| 7. Thyrotropes | Tsh | 59 | 2.3 | 46 | 1.2 | *tshba* | <sup>14,15</sup> |
| 8. Fsh- gonadotropes | Fsh | 99 | 3.8 | 198 | 5.2 | *fshb* | <sup>15,32</sup> |
| 9. Lh-gonadotropes | Lh | 416 | 16.1 | 895 | 23.5 | *lhb* | <sup>2,15,21,31</sup> |
| 10. Gonadotrope-like |  | 668 | 25.8 | 444 | 11.7 | *cga, nr5a1b* | <sup>32,39</sup> |
| 11. Red blood cells |  | 364 | 14.0 | 559 | 14.7 | *gb* |  |
| 12. Macrophages |  | 15 | 0.6 | 15 | 0.4 | *c1qa, c1qb, c1qc* | <sup>14,15</sup> |
| 13. Uncharacterized |  | 299 | 11.5 | 410 | 10.8 |  |  |
| 14. Uncharacterized |  | 197 | 7.6 | 386 | 10.2 |  |  |
| 15. *cga*-expressing cells |  | 115 | 4.4 | 160 | 4.2 | *cga* | <sup>32</sup> |
| 16. Uncharacterized |  |  |  | 21 | 0.6 |  |  |

### Cluster refinement

During initial data exploration, we observed several clusters sharing many common markers. Therefore, we further inspected their heterogeneity iterating the clustering function on selected subsets of the data. Sub-clusters were detected across multiple clustering resolutions (0.2 and 0.5) of the *FindClusters* function in Seurat. We subsequently used the *FindAllMarkers* function to find differentially expressed genes between each of these sub-clusters to highlight the differences and finally decided on the cell-type boundaries for these ‘zoomed-in’ sub-clusters.

## Data Records

We present a characterization of the cell types of the medaka pituitary gland, as a basis for more in-depths analyses of the regulation of and inter-communication between different hormones and cell types.

Our data set on the medaka pituitary consists of gene expression values for 2592 and 3804 adult female and male cells, respectively. These can be assigned to 15 or 16 distinct cell populations, respectively, one of which is unique to male pituitaries. We assigned biological identities to 12 populations based on the expression of known hormone-encoding genes. Fig. 1 and Table 4 provide an overview of the cell types of the medaka pituitary.

**Figure 1.**
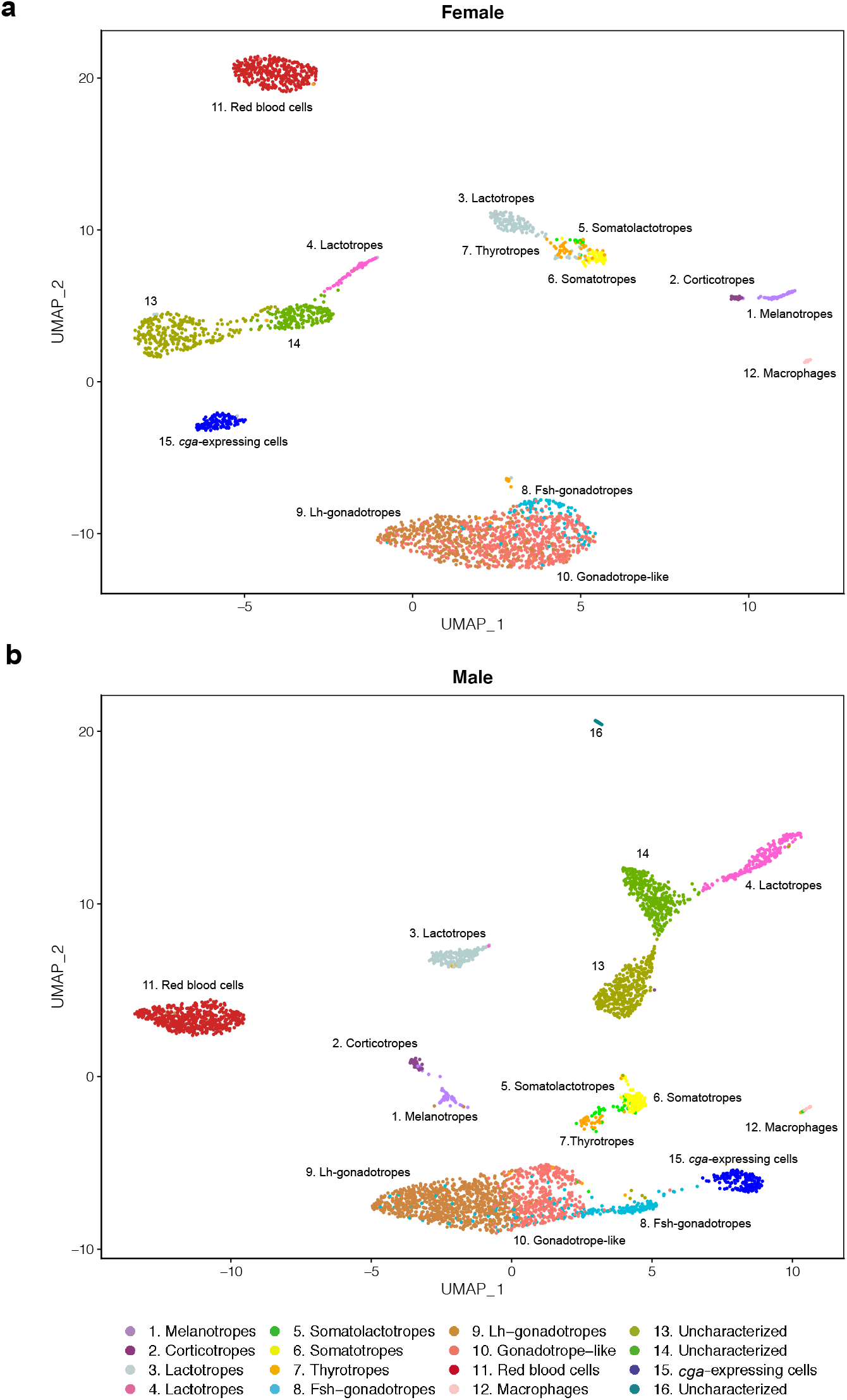
Clustering of scRNA-seq data reveals the cell populations of the female and male medaka pituitary. (a) Uniform manifold approximation and projection (UMAP) plot showing the classification of distinct cell types of the female pituitary (b) UMAP plot showing the classification of distinct cell types of the male pituitary. UMAP offers a projection of individual cellular expression profiles (which are high-dimensional data points) into two dimensions. Cells are represented by dots; close proximity of cells in the UMAP projection indicates their expression profiles are similar. Therefore, cells of a distinct type appear as clusters. See Table 4 for details on cluster identity assignment.

### Availability

The main data record consists of *barcodes.tsv, features.tsv* and *matrix.mtx* files, listing raw UMI counts for each gene (feature) in each cell (barcode) in a sparse matrix format. In addition, we provide a table of cell type assignments and UMAP projections for each individual cell (barcode), as well as a summary of the top differentially expressed genes per cell type. Raw sequencing data are available in FASTQ format. Finally, the data record contains matrix files on exonic and intronic expression, which can be used for Velocyto gene expression dynamics analyses^33^. These data can be accessed through the project accession number GSE162787 at the NCBI Gene Expression Omnibus (GEO: *https://ncbi.nlm.nih.gov/geo/*).

## Technical Validation

In order to validate the quality of our data, we investigated both technical variables and the biological perspective.

### Technical quality control

Interpretation of single-cell transcriptomics data is highly sensitive to technical artifacts. For example, expression profiles can derive from low quality cells, cellular debris, or ambient RNA. We therefore determined empirical thresholds distinguishing genuine cellular expression profiles from noise (Fig. 2). We defined selection criteria for cellular barcodes based on UMIs counts (Fig. 2a), expression of mitochondrial genes (Fig. 2b, cyan dots), expression of genes specific to red blood cells (Fig. 2c, red dots), and a probability score for cellular doublets (Fig. 2, orange dots). Based on the distribution of UMI counts per barcode, we defined as cellular barcodes those with ≥1500 UMIs in both female and male samples. High levels of expression of mitochondrial genes have been interpreted as an indication of damaged cells^34,35^.We therefore excluded barcodes in which mitochondrial gene expression contributes more than 5% of the total (for both females and males, Fig. 2b).

**Figure 2.**
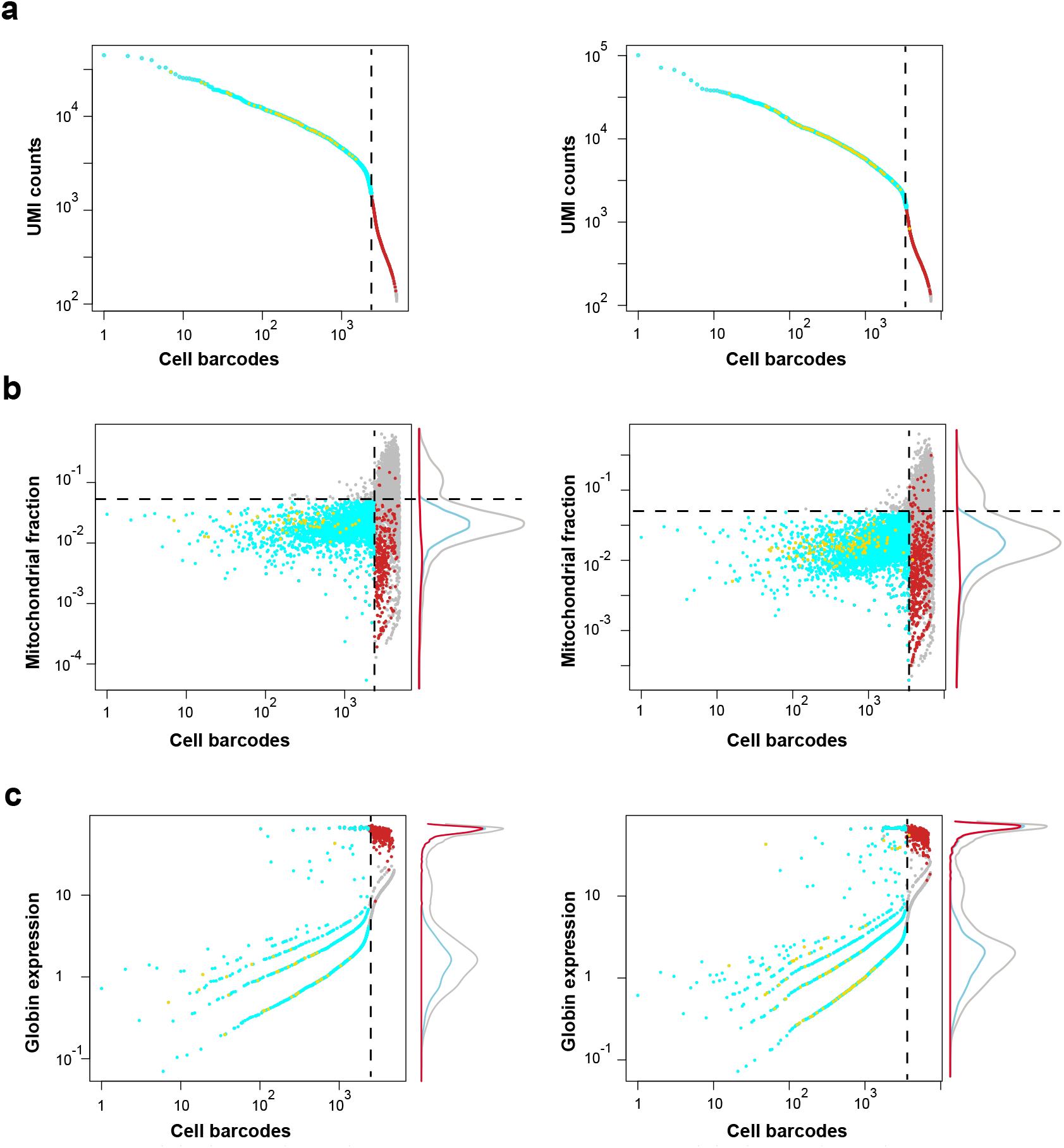
Selection of cellular barcodes in pituitary single-cell data. (a) Threshold 1: QC based on the UMIs counts. The cyan dots illustrate the final selected barcodes, grey dots represent barcodes not meeting selection criteria, orange dots indicate potential doublets, and red dots represent globin-expressing cells. The dotted line represents the UMI cutoff value. (b) Threshold 2: QC based on the mitochondrial fraction. The horizontal dotted line indicates the 5% threshold. Marginal plots indicate the densities of barcode sets at different values (grey line: all cells). (c) Threshold 3: QC based on globin (*gb*)-expressing cells (red dots), which do not meet the UMI count criterion (panel and dotted line) but are included in our final dataset. Note that the axes are on logarithmic scales, and that the horizontal axis is identical in all panels.

In initial clustering analyses, we noticed that these criteria exclude both technical artifacts (data not shown) and a single cluster of cells with low overall UMI counts, but a very clearly defined expression profile. Based on their expression profile (Fig. 3b) we interpret these barcodes to represent red blood cells. Although they are developmentally only very distantly related to the other cells in the pituitary, our data show that they are an intrinsic (if transient) component. Therefore, including them in our dataset will allow for more precise interpretation of patterns in bulk RNA-seq studies.

**Figure 3.**
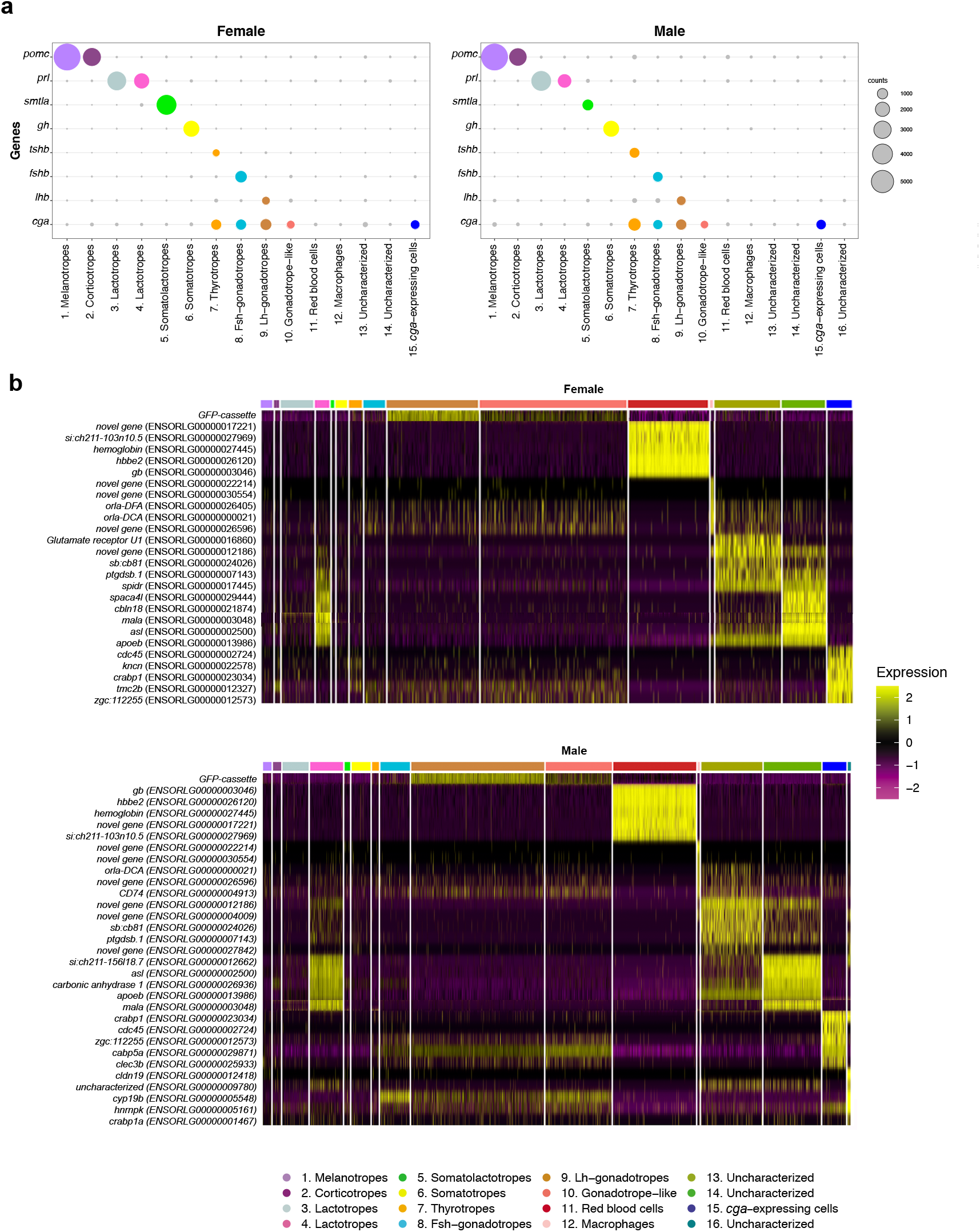
Cell type functions. (a) The expression of hormone-encoding genes in female and male pituitary, respectively. Values are the mean of normalized expression over all cells in a cluster. The grey dots indicate low expression of genes. (b) Expression pattern heatmaps for uncharacterized cell types in both female and male data.

### Biological quality control

In order to validate the biological relevance of the cell type identities shown in Fig. 1, we have compared their gene expression patterns with known expression profiles from both teleosts and mammals. Initially, we identified eleven different populations based on optimized Seurat clustering. However, there were several clusters that shared high expression of known, specific marker genes. We therefore evaluated expression heterogeneity within each cluster and manually defined final clusters (see Methods). We defined two subclusters for the initial cluster expressing *pomca* (ENS0RLG00000025908), which encodes pro-opiomelanocortin, the protein precursor for α-Msh and Acth). The first subcluster has high expression of *pomca, oacyl* (ENS0RLG00000005993), *crhbp* (ENS0RLG00000002228), *pcsk2* (ENS0RLG00000006472) and *pax7* (ENS0RLG00000004269), which are markers for the α-Msh-secreting melanotropes; the second subcluster was identified as Acth-secreting corticotropes due to lower expression of *pax7* and *pomca*^15^.

Next, a large compound cluster on the right side of Fig. 1 (both females and males) shares three important hormone-encoding genes: *tshb, gh* and *smtla*. We therefore divided it into three subclusters using Seurat, gene expression patterns and differentially expressed (DE) genes (see available code for details). The first subcluster has strong expression of *tshba* (ENS0RLG00000029251, encoding the β subunit of Tsh), which is a robust marker for thyrotropes (orange color in Fig. 1). The second subcluster expresses *gh* (ENS0RLG00000019556), which demarcates the somatotropes (light green). The last subcluster (yellow) has expression of *smtla* (ENS0RLG00000013460, which encodes Sl, a hormone produced by the teleost, but not the mammalian, pituitary gland)^16,21,31,36^. In contrast to in the zebrafish pituitary^16^, we did not observe the expression of *pou1f1* (ENS0RLG00000015870) in somatolactotropes.

Finally, Seurat initially divided the big cluster of gonadotrope cells (expressing the gonadotropin α-subunit *cga*, ENS0RLG00000022598) at the bottom of Fig. 1 into two subclusters for the males, but not for the females. However, based on the heterogeneity of the expression pattern in the female cluster, and its apparent homology to the two male clusters, we defined two subclusters: one is enriched for *lhb* (ENS0RLG00000003553, encoding the β subunit of Lh) and *gnrhr2* (ENS0RLG00000012659, also known as *gnrhr2a^37^*) expression, both of which are exclusively associated with Lh production in teleosts^37,38^; the other expresses *nr5a1b* (ENS0RLG00000013196), encoding a nuclear receptor involved in gonadotropin regulation in the mammalian pituitary and teleosts^16,39^.

Three remaining cell clusters we directly assigned to specific hormone-producing cell types based on the very high expression of hormone-encoding genes: two distinct populations of lactotropes expressing *prl* (ENS0RLG00000016928) and *fshb*-expressing gonadotropes (ENS0RLG00000029237, encoding the β subunit of Fsh).

After final clustering, we define 15 and 16 distinct expression patterns in female and male data, respectively, which we interpret as distinct cell types (see Fig. 1 and Table 4). Based on their expression patterns, we could assign unambiguous cell type identities to twelve of these (Fig. 3). Ten clusters show high expression of genes encoding protein hormones (Fig. 3a and Table 4). In addition, one cluster expresses *cga*, but no gene encoding a β subunit. Two clusters appear to be red blood cells and macrophages (Fig. 3b and Table 4). Based on detailed expression patterns (Fig. 3b) we could not unequivocally define the elusive folliculostellate cell population of the pituitary. In the mammalian pituitary, these cells (of partly unknown function) are characterized by marker genes^14^ that show significant, but specific expression in two of our uncharacterized cell populations (clusters 13 and 14, Fig. 1).

Traditionally, a marker for these cells is the *s100* gene^40,41^ however, this gene has at least eight homologs in the medaka genome, none of which shows a clear expression pattern in the pituitary.

Overall, our assignments of biological function largely confirm a ‘one cell type, one hormone’ (Fig. 3a) division of labour, in which major hormones are produced by a single, dedicated cell type. This is in apparent contrast to the mammalian pituitary, in which cells are often responsible for producing and releasing multiple hormones^14^.

## Code Availability

The R code used in the analysis of the scRNA-seq data is available on GitHub (https://github.com/sikh09/Medaka-pituitary-scRNA-seq).

## Acknowledgements

We are thankful to Susanne Lorenz for sequencing and Lourdes Carreon G Tan for fish facility maintenance. This work was supported by NMBU and the Norwegian Research Council grants no 251307, 255601, and 248828.

## Author contributions

Eirill Ager-Wick, Finn-Arne Weltzien and Christiaan Henkel conceived and designed the project. Eirill Ager-Wick isolated the cells. Khadeeja Siddique, Romain Fontaine and Christiaan Henkel analyzed the data and wrote the paper with input from all other authors.

